# Mathematical Modeling of Neural Stem Cell Migration within Brain using Multi-Fiber Tractography

**DOI:** 10.1101/2025.02.13.638130

**Authors:** Austin Hansen, Russell Rockne, Vikram Adhikarla, Margarita Gutova, Heyrim Cho

## Abstract

An agent-based model of therapeutic neural stem cell (NSC) migration is developed and used to predict the migration of NSCs in naïve mouse brain. The model utilizes generalized q-sampling imaging which resolves white matter fibers that cross in the brain and is shown to better account for variations in NSC migration patterns as compared to diffusion tensor imaging. In calibrating the model to experimental data, we show that the model is able to reproduce the distribution of NSCs in the mouse brain. In addition, we show that the distribution of NSCs in the mouse brain is sensitive to the location of NSC injection. Persistent distribution of NSCs to the olfactory bulb, consistent with developmental pathways including the rostral migratory stream, suggests that future models of therapeutic NSCs in the naïve brain may need to include other factors such as chemotaxis or blood flow to account for variations in NSC migration paths. The results highlight the usefulness of the model in predicting which injection locations may provide the best distribution of NSCs to a given target location.

## 1 Introduction

Neurological disorders encompass a wide range of different ailments and are estimated to be one of the top contributors to the overall disease burden [42]. Despite their prevalence, current treatments for many of these disorders, including Alzheimer’s, Parkinson’s, Huntington’s, multiple sclerosis, glioblastoma, and traumatic brain injuries, are only able to treat symptoms and are unable to address fundamental drivers. For this reason, neural stem cell (NSC) therapies have emerged as a promising direction [27]. Unlike many other treatments, neural stem cells are able to address multiple mechanisms of action ranging from cell replacement to immunomodulation, and can even be engineered to carry therapeutic agents to the injury sites [36, 49, 19, 6, 21, 53]. Because of the broad array of benefits, many clinical trials have been completed or are currently being conducted to test the safety and efficacy of these treatments [8, 39, 46, 37, 26, 38, 18, 4]. In some cases, treatments such as autologous hematopoietic stem cell transplantation have become more routinely used for patients with multiple sclerosis [30, 34].

One of the hurdles that must be addressed on the path to clinical implementation of many NSC treatments, however, is optimizing delivery strategies to ensure that an adequate number of viable cells reach the target site. Since in vivo experiments can be costly and time-consuming, mathematical modeling can provide a platform to test delivery strategies and inform both experimental design as well as clinical recommendations. For this reason, mathematical modeling has been used to great effect in understanding some neurological disorders, as well as tumor growth, and more particularly cancer cell migration and invasion [31, 3, 35, 28, 32]. For reviews of various modeling techniques, we refer the reader to the following [11, 5, 10, 50, 12]. In [40] a partial differential model was used to study the migration of neural crest cells (NCCs) in the gut and found that the proliferation of a small number of cells at the leading edge was the main contributing force to their migration. Other continuous approaches have tried to include nonlocal interactions such as cell-cell contact inhibition or attraction within continuum approaches by means of integro-partial differential equations as seen in [33]. Despite these successes, continuum models are often unable to capture more detailed cellular characteristics. For this reason, both individual-based and hybrid models have been employed. In [29], an agent-based model confirmed that efficient streaming of NCCs relied on the presence of two distinct phenotypes, with only a few cells near the front taking the role of leader cells and the rest being followers. In [15] the utilization of a random walk approach allowed the authors to account for the effects of the Rac1 enzyme on the motion of neural crest cells, while still providing the ability to derive continuous approximations of NCC migration. Despite several works related to NCCs, very few models have been developed for the migration of therapeutic neural stem cells.

Of the few models of neural stem cell migration in cancer therapy, both have been primarily based on tissue anisotropy [38, 17]. This is in part due to the fact that many cells, including NSCs, show preferential movement along white matter tracts. Although the exact mechanism for this behavior is largely unknown, many studies have begun to take tissue structure into account as a force in cell migration. For example, computational models of glial cells which include the effects of tissue anisotropy in addition to diffusion and proliferation, are better able to capture the geometry of glioma than those that exclude tissue anisotropy [43]. In addition, NSC clusters were shown to be oriented along the directions of white matter fibers [38]. However, these models rely on diffusion tensor imaging (DTI) which is subject to a number of challenges stemming from its inability to resolve multiple crossing fibers within a single voxel. Specifically, fractional anisotropy (FA) is unable to accurately capture specific microstructures in places of crossing fibers and is also insensitive to certain changes in white matter that come from damage or disease, which may be crucial in pathologies like glioma and traumatic brain injury.

For example, it is well known that axional diffusivity and radial diffusivity, both of which come from DTI, are often used to measure axonal and myelin damage. However, these measures can be affected by swelling and edema in ways that more advanced imaging techniques such as q-space imaging are not [51]. In one study, it was shown that for mild traumatic brain injury, DTI was not as sensitive as q-space analyses to changes in axonal volume, which is common during axonal shearing or twisting, among other causes [48]. For a good review of the advantages and disadvantages of different imaging techniques, see [51]. Despite challenges with DTI, generalized Q-ball imaging (GQI) has been shown to accurately capture multiple crossing fibers in a single voxel, and was subject to less artifacts like deletions or incompletion when compared to DTI [24]. Moreover, in a study that examined 18 patients with gliomas, it was found that Q-ball imaging (QBI) was better able to capture metrics such as tract volume, density, and length [7]. GQI has also been suggested to aid in preoperative planning for patients with low-grade gliomas by helping to distinguish areas of the brain in which the glioma has infiltrated or displaced certain white matter tracts [13].

Thus, in order for NSC migration models to be adequately predictive for use in clinical settings, the utilization of more robust imaging techniques is a critical step. To address these limitations, we build upon the model in [38, 17] in two ways: first through the use of GQI reconstruction we are able to create a probabilistic model which captures competing crossing fibers [14, 52, 24]; second we alter the model to allow each cell’s motion to be a continuous sum of directed and random motion based on the underlying anisotropy. Section 2 outlines the model equations and parameters, section 3 explains the experimental data to which the model is then calibrated to, while section 4 outlines the results from our parameter study, as well as the calibration and validation. Finally, in section 5 we discuss some of the limitations of the model and future directions for this work.

## 2 Mathematical model of neural stem cell migration

We assume tissue structure is the primary driving force of neural stem cell migration in the absence of chemical signals. Furthermore, since white matter tracts are highly aligned while grey matter is less structured, the direction of a neural stem at a given time point will be a combination of directed motion along the white matter tract and random motion, with the relative proportion dependent on the degree of anisotropy [44]. Thus, the equation for the position of a neural stem cell ***x***_*k*+1_ given its position at time *k* is as follows:

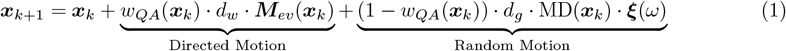

where, *w*_*QA*_(***x***_*k*_) is a scaling factor proportional to the degree of anisotropy an position ***x***_*k*_, *d*_*w*_ is the maximum step size along white matter, *d*_*g*_ is the maximum step size along grey matter, ***M***_*ev*_(***x***_*k*_) is the white matter fiber direction, MD(***x***_*k*_) is the mean diffusivity, and ***ξ***(*ω*) is a random sample on a 3D unit sphere.

The direction of the vector ***M***_*eυ*_ is probabilistically chosen based on the first three fiber directions resolved through GQI. In GQI reconstruction, each fiber direction is given a quantitative anisotropy (QA) value, which measures the increased diffusion along a given direction over the background isotropic diffusion. The higher the QA value in a direction, the stronger the tissue alignment in this direction. We use GQI in conjunction with traditional DTI reconstruction to build a model that can accurately differentiate the following scenarios regarding fiber alignment and fiber density shown in Fig. 1, denoted as cases (A–D).

**Figure 1.**
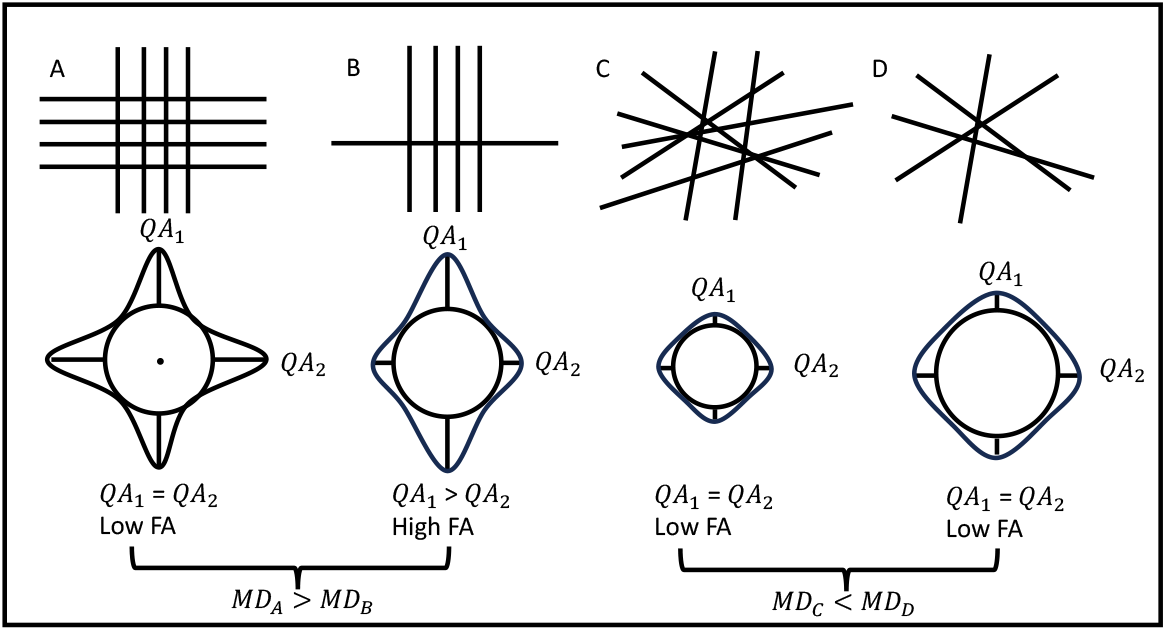
Different scenarios of tissue orientations in the top row and the corresponding resolution from GQI in the bottom row. (A) Four equal crossing fibers which are highly aligned, (B) a single white matter fiber crossing 4 aligned fibers, (C) densely packed region with little to no alignment of any fibers, (D) a sparsely packed region with little to no alignment of any fibers. Quantitative anisotropy values (*QA*_1_, *QA*_2_), fractional anisotropy (FA), and mean diffusivity (MD).

**Figure 2.**
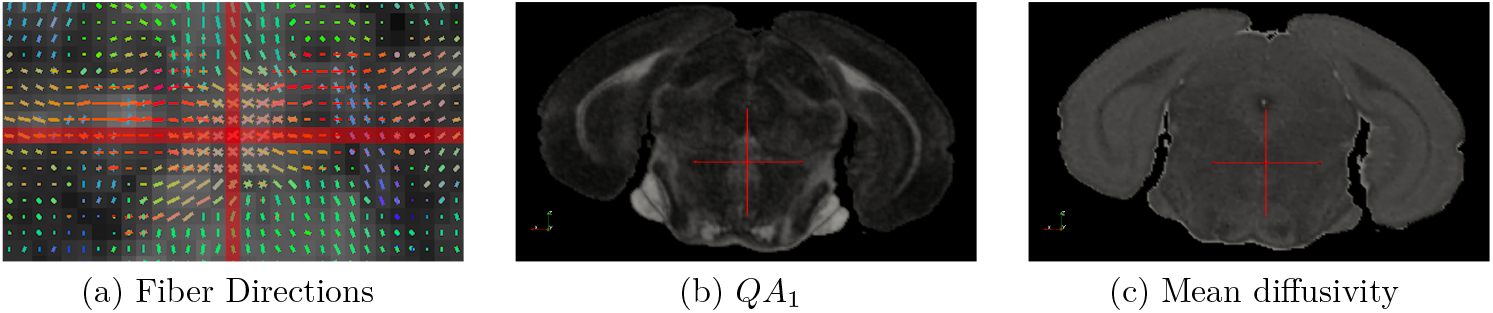
(a) Fiber directions as visualized in DSI-Studio [1] for a given cross section. (b) Cross section of the of the first quantitative anisotropy value (*QA*_1_). (c) Cross section of the of the mean diffusivity.

In the case of several highly aligned fibers crossing, or a single fiber crossing several aligned fibers (Fig. 1 A and B), GQI resolves both multiple crossing fibers, as well as locations with only one major fiber direction. For example, for Fig. 1A, GQI would yield two large QA values, *QA*_1_ and *QA*_2_, for each of the major fiber directions. In contrast, B depicts a scenario in which *QA*_1_ > *QA*_2_ representing only one major fiber at this location.

For modeling purposes we only consider the first three QA values obtained from GQI reconstruction, denoted as *QA*_1_, *QA*_2_, and *QA*_3_, corresponding to the fiber directions, dir_1_, dir_2_, and dir_3_, respectively. Furthermore, in order to reduce the noise from any insignificant non-primary fiber directions, we eliminate secondary and tertiary QA values that are less than 0.1 by making them zero. Thus, when a NSC is at a location with multiple QA values greater than 0.1, the migration direction is probabilistically chosen based on both the relative QA values and the momentum of the cell. Here we include some momentum within the direction selection so as to allow a NSC a greater ability to follow a secondary/tertiary fiber once it has been chosen. The probability of following the direction of the *i*-th fiber based on anisotropy is defined as

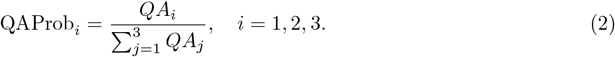

To model the probability of choosing the *i*-th fiber due to momentum we compute the angle between the NSC’s motion in the previous time step, NSC_dir_ = ***x***_*k*_−_1_ − **x**_k_−_2_, and the *i*-th fiber direction dir_*i*_. Suppose *n*_*i*_ is the number of fiber directions at position ***x***_*k*_.

Then we have

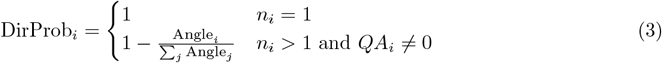

Since it is unknown how much either of these factors contribute the to motion independently, we assume that the overall probability is equally weighted between both. Hence the final equation for the probability of choosing the *i*-th fiber is

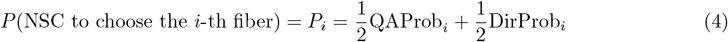

The direction ***M***_*eυ*_ = dir_*i*_ is then selected at each time step based on the calculated probability for each cell at its current location.

In the case where there is little tissue alignment, quantitative anisotropy alone cannot distinguish those regions of the brain in which tissues are tightly packed from those in which there is less density due to factors such as myelin degradation or cerebrospinal fluid. In order to help distinguish between these scenarios (Fig. 1 C, D), we use mean diffusivity as a measure of overall diffusion. The mean diffusivity MD is defined as the average of the three eigenvectors resolved through DTI reconstruction, and is typically measured in 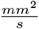. Here we only consider the effects of mean diffusivity within the randomized movement. This is done by multiplying the grey matter velocity by 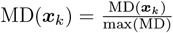, where MD is the mean diffusivity. Thus, the higher the mean diffusivity, the more random motion will contribute to the overall motion. Note that this also has the effect of making the contribution from directed motion larger when both step sizes *d*_*g*_ and *d*_*w*_ are equal.

Both parameters *d*_*w*_ and *d*_*g*_ represent the step size that a neural stem cell can travel in a given time step along the white matter and grey matter, respectively. Due to heterogeneity in neural stem cell populations, not all cells have the same motility. We account for this stochasticity by modeling *d*_*g*_ and *d*_*w*_ as

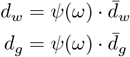

where *ψ*(*ω*) ∼ B[*α, β*] is a random variable with beta distribution, and 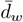 and 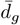 are the maximum step sizes on white and grey matter, respectively. Furthermore, we take the parameters in the beta distribution as *α, β* ≥ 1. Note when *β* > 1 and *α* = 1 the majority of cells move with a slower velocity, while only a few outliers move more quickly. In contrast, when *α* > 1 and *β* = 1, most cells have a higher velocity and only a few move comparatively slow.

The overall proportion of directed vs. random motion for a cell at position ***x*** is determined by the *QA* value of the chosen direction ***M***_*eυ*_. Thus, *w*_*QA*_ is represented by the hill function

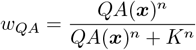

Here the closer *w*_*QA*_ is to 1 the more of its motion is directed by the tissue structure. In contrast, the closer *w*_*QA*_ is to 0, the more random the motion becomes. In addition, we point out that *K* represents the QA value in which half of the motion comes from the fiber direction and the other half is a random walk.

## 3 Experimental Data

In order to calibrate the model we use time series data taken from previously published results examining the migration of NSCs in the naïve mouse brain [38]. The NSCs are developed from allogeneic human NSCs and genetically modified to express human L-myc gene (LMNSC01) [27]. These NSCs demonstrate efficient cell migration within brain directed by brain tissue and endogenous cues and have promising therapeutic potential because they also inherently migrate to sites of central nervous system damage, such as brain tumors and brain injuries [20, 6, 54, 21, 19]. Moreover, the NSCs can differentiate into neurons and glial cells in vitro and in vivo and can contribute to recovery from brain damages via cell replacement and neuroprotective effects [27, 38, 4, 18].

In [38], the authors inject the NSCs, namely, LMNSC01 cells near the right frontal corpus callosum and then euthanized mice at 3,6, and 9 months. Brain tissue was stained for NSCs for each mouse and the median distance from the injection site, as well as the percentage of cells in white matter was quantified for each of the 3 groups with 5 mice in each group. Additional details of the experimental design are provided in [38].

The median distance from the injection site was 437*µm*, 1030*µm*, and 902*µm* at 3, 6, and 9 months respectively, while the percentage in white matter was 66%, 72%, and 59%. In addition, the NSC’s were largely localized in the corpus callosum at 3 months, migrated to the anterior commissure by 6 months, and to the olfactory bulb by 9 months.

Using 2D tissue slices of the brain volumes at 6 and 9 months, we quantified the percent of injected NSCs in the anterior commissure and olfactory bulb. Quantification was performed with the Color Thresholder tool in MATLAB. The original images and segmentations are shown in Figure 3. We estimated that there are an average of 7.2% of cells in the anterior commissure by 6 months, 8.5% of cells in the anterior commissure by 9 months, and 19.3% of cells in the olfactory bulb by 9 months.

**Figure 3.**
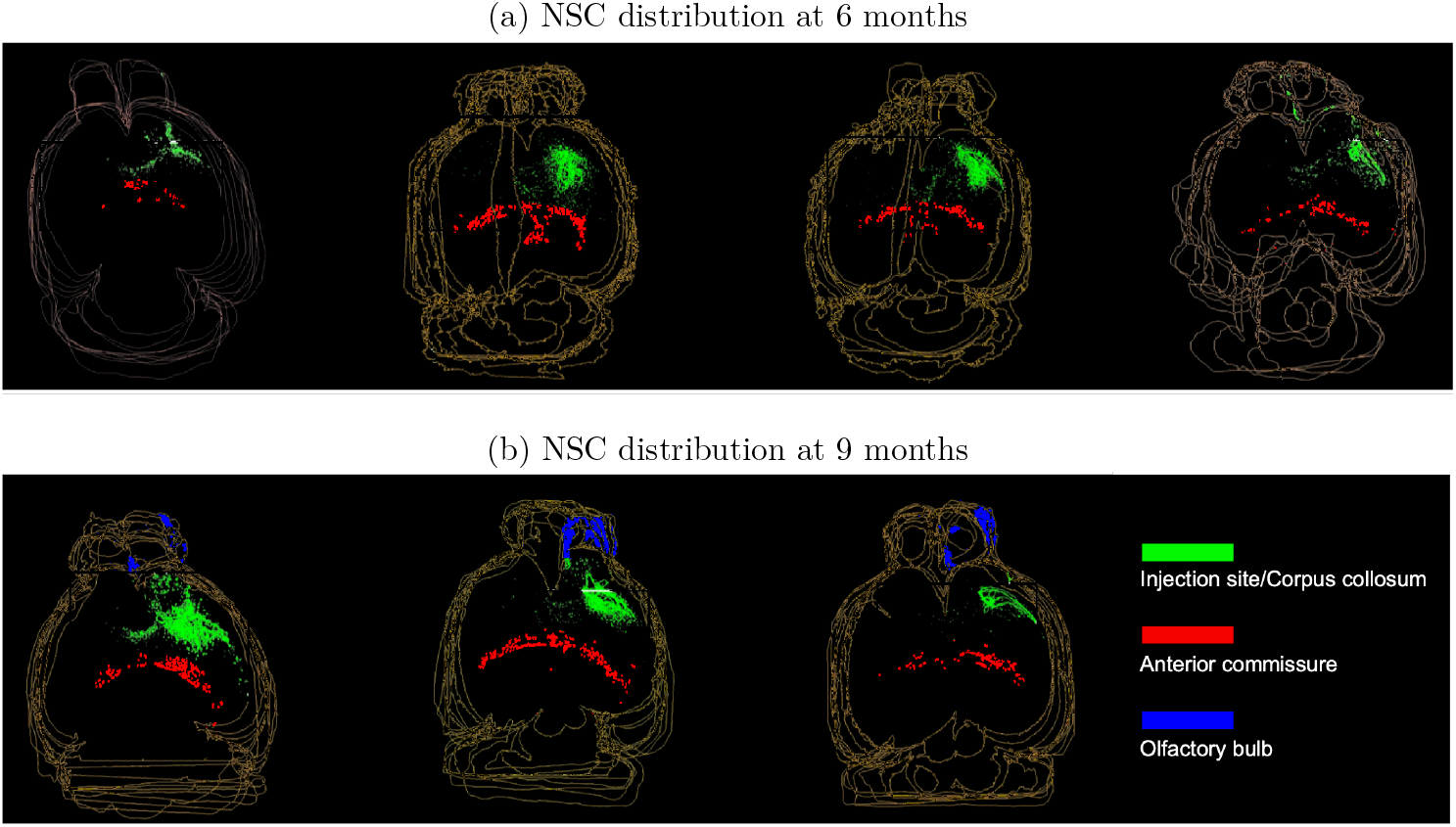
Three dimensional analysis of NSC distribution created from sequential 2D tissue sections from mice euthanized at (a) 6 months and (b) 9 months. NSCs are segmented and shown in the anterior commissure, the olfactory bulb, and the injection site/corpus callosum.

## 4 Results

### 4.1 Comparison between Single and Multi-Fiber Migration Models

We first illustrate the difference between a single fiber migration model using DTI and the probabilistic multi-fiber migration model based on GQI. For the single fiber migration model, the equation of motion is the same as in equation 1, except *M*_*eυ*_ is set to the single direction resolved from DTI and *QA*_1_ = FA while *QA*_2_, *QA*_3_ = 0. Here FA is fractional anisotropy calculated from the eigenvalues of the diffusion tensor. To accurately isolate the differences from the addition of multiple fibers, we remove the stochasticity parameters from the step sizes and set *d*_*g*_ = 0.1 and *d*_*w*_ = 0.1. The simulation is initialized with 100 NSCs in a single voxel located within a region with a high density of secondary fibers, shown in Figure 4 as a black dot. The simulations are run for 100 time steps equating to 1 day. Other parameters are set as *K* = 0.18, *n* = 7, and *α* = *β* = 1.

**Figure 4.**
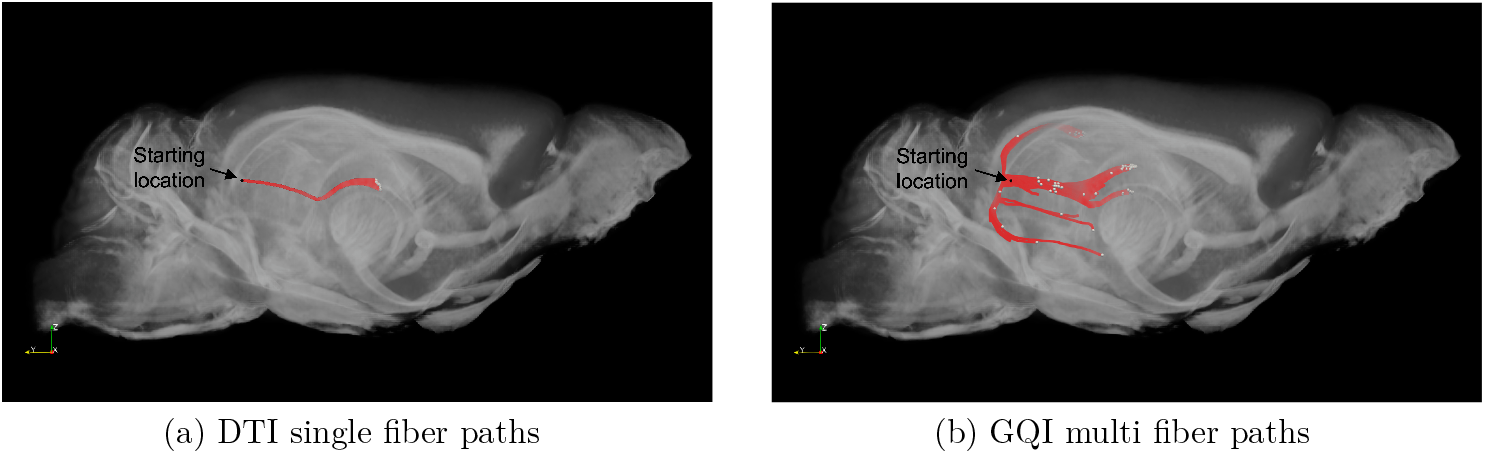
Comparison of simulated NSC migration trajectories. (a) 100 trajectories using DTI, which only resolves the primary fiber direction. (b) 100 trajectories using GQI reconstruction, which resolves multiple crossing fibers.

In Figure 4(a) we see the migratory pathways from the primary fiber model based on DTI data. Here we see that all 100 cells roughly follow the same path and only near the very end do the cells start to show some hints of diverging into different trajectories. In contrast, the probabilistic multi-fiber model based on GQI reconstruction in Figure 4(b) shows multiple different pathways accessible to the cells. We can see that some cells follow the same white matter fiber as before, but we also have some secondary and tertiary fibers that lead vertically. This allows the cells to move up and down before finding fibers that are directed toward the front of the brain. The results of the multi-fiber model demonstrate the NSCs’ enhanced ability to explore a broader range of regions within the brain.

### 4.2 Parameter Sensitivity Analysis

To study the impact of model parameters on the predictions, we performed a sensitivity analysis on the model parameters 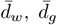, *K, n, α*, and *β*. To do this, we performed 400 simulations in which parameter values were chosen with Latin Hypercube Sampling (LHS) [23, 22] with 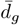 and 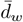 ∈ [0, .6], *K* ∈ [0, .5], *n* ∈ [1, 30] and *α, β* ∈ [1, 100]. The partial rank correlation coefficients (PRCC) were then calculated between each parameter and the median distance from injection site and percent of NSCs in white matter at 9 months (Figure 5).

**Figure 5.**
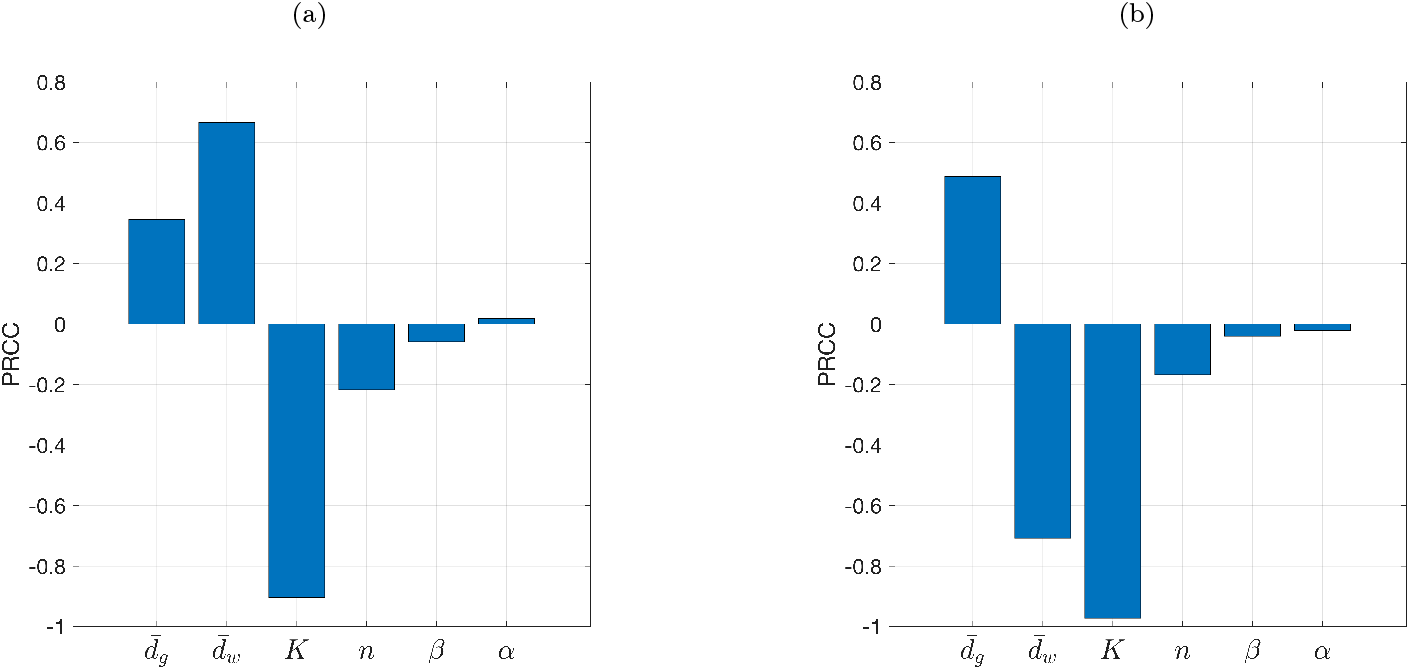
Partial rank correlation coefficients (PRCC) for 6 model parameters and their effect on (a) median distance from injection site and (b) percentage of NSC in white matter at 9 months.

We find that 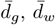, and *K* had the largest effect on the median distance from the injection site, as well as the percent of NSCs in white matter at 9 months. In particular, we find that the level of anisotropy in which motion is half directed and half random (*K*) is inversely related to the median distance and percent in white matter, while the Hill coefficient *n*, and stochastic migration speed parameters *α* and *β* do not have a significant impact on either of these quantities. However, we note that *α* and *β* govern the number of outliers in the distance from the injection site despite not having a large overall effect on the median distances.

In addition, we find that the white matter step size, 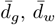, is positively correlated with the median distances and is inversely related to the percent in white matter, while the grey matter step size, 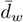, is positively related to both the median distance and the percent in white matter. However, we do observe some non-monotonic behavior for the parameter 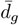 when we plot the residuals 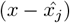 and (*y*_*j*_ − ŷ) where *x* and *y* are the rank transformed 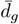 values and 9 month median distances respectively, and 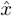, *ŷ* are from the linear regression models. The residual for 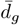 is depicted in Figure S2 of the supplementary file. To identify the cause, we examined the parameter values that corresponded to these outliers and found that the majority of the outliers occurred with either a very low 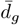 or 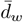 value. Given this information, we calculated the PRCC for a set of simulations in which 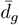 and 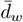 were varied evenly in the range [.0025, .15] and all other parameters were held constant with *K* = .25, *n* = 7, and *α, β* = 1. In this we find that the PRCC value for 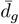 and the corresponding residual plot shows a strong negatively correlated relationship.

In addition to this change in PRCC for low step sizes, we also hypothesize that the ratio of 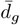 and 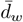 may play a crucial role given their opposite effects on the percentage of white matter. In order to explore this, we look at how the change in percent of NSCs in white matter varies with the ratio of 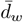 to 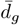. In Figure 6(a), we plot the change in the percentage of NSCs in white matter measured by Δ*W* = *W* (*t*_end_) − W (t_0_), where *W* (*t*) is the percentage of NSCs in white matter at time *t*. Here, *t*_0_ is the initial time and *t*_end_ = 27000 or 9 months time. We see that the change in the percentage on white mater is positive when 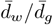 is very small, but as the ratio 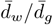 increases, the change in percentage becomes negative and decreases rapidly. In other words, when 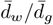 is large, NSCs tend to move out of the white matter. This implies that when the random movement, 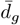, dominates the migration, the cells collect in white matter, and when directed movement, 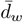, dominates the migration, the cells tend to move to more grey matter locations. This depicts the inverse relation between the change in percent in white matter Δ*W* and 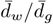, and that Δ*W* may be an important measure in calibrating the step size parameters.

**Figure 6.**
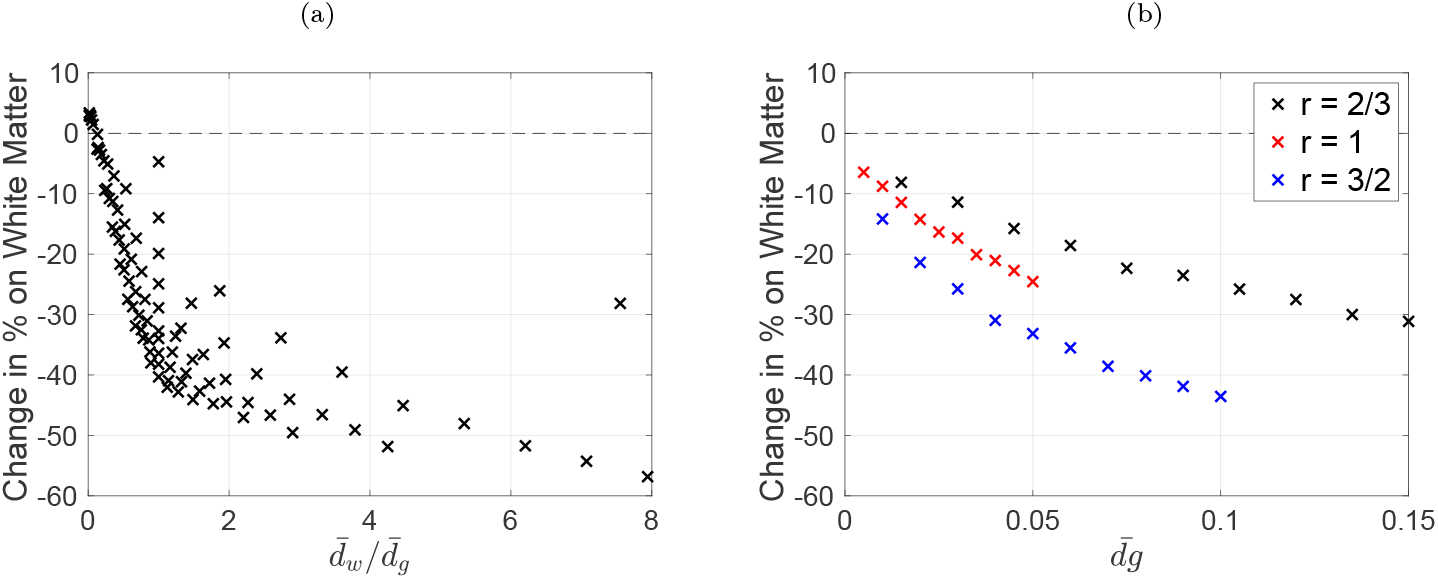
Change of percentage of NSCs in white matter depending on migration step sizes, 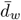 and 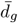. (a) The percent of NSCs in white matter decreases rapidly as the ratio of directed and random movement step sizes, 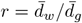, increases. (b) The percent change in white matter is plotted for fixed step size ratios, *r* = 2*/*3, 1, 3*/*2. The percent in white matter also decays as the magnitude of step size 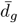 increases.

This, however, does not show the whole picture as we can see for each ratio there is some variance in the change in percentage in white matter. Thus, we consider three different fixed ratios 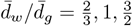, and observe how the change in percentage in white matter varies as the magnitude of the step sizes increases. To accomplish this, we set 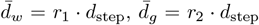 and vary *d*_step_ ∈ [.005 : .05] for a fixed 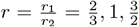. Ten simulations were performed for each ratio. Figure 6(b) shows that as the magnitude of the step sizes increases, the cells slowly occupy more grey matter. Thus not only does the ratio matter, but the velocity of the cells helps to determine how many cells are on white or grey matter by the end of the simulations.

### 4.3 Injection Location and Path Clustering Analysis Toward Target Regions

In this section, we study the effects of injection location on the migration paths to specific target regions. An important qualitative trend found in the experiments reported in section 3 is the accumulation of NSCs in the anterior commissure at 6 months and the olfactory bulb at 9 months. We observed that in addition to the model parameters, the injection location had a significant effect in determining the location of NSCs in the brain. Furthermore, since we estimated the percentage of NSCs in the olfactory bulb to be ≈ 19% at 9 months, we hypothesized that cells must travel along white matter tracts to reach this region in sufficient quantities.

To identify potential pathways to the anterior commissure and olfactory bulb from the injection site, we simulated an injection of 524, 536 cells into the corpus callosum and simulated their migration for 9 months. The number of cells was chosen so that four cells were initialized at each lattice point within the corpus callosum. This was done for nine combinations of 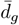 and 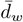 values, with each being selected from the following set {0.1, 0.35, 0.6}. Figure 7 shows a heat maps of the NSC injection locations that resulted in NSCs migrating to the anterior commissure or the olfactory bulb at any point during the simulation. The plotted quantity shows the number of NSCs that started at the corresponding injection location. We note that the plotted values are the maximum value of all the vertical *Z* slices within each *X/Y* bin, to visualize it from top view.

**Figure 7.**
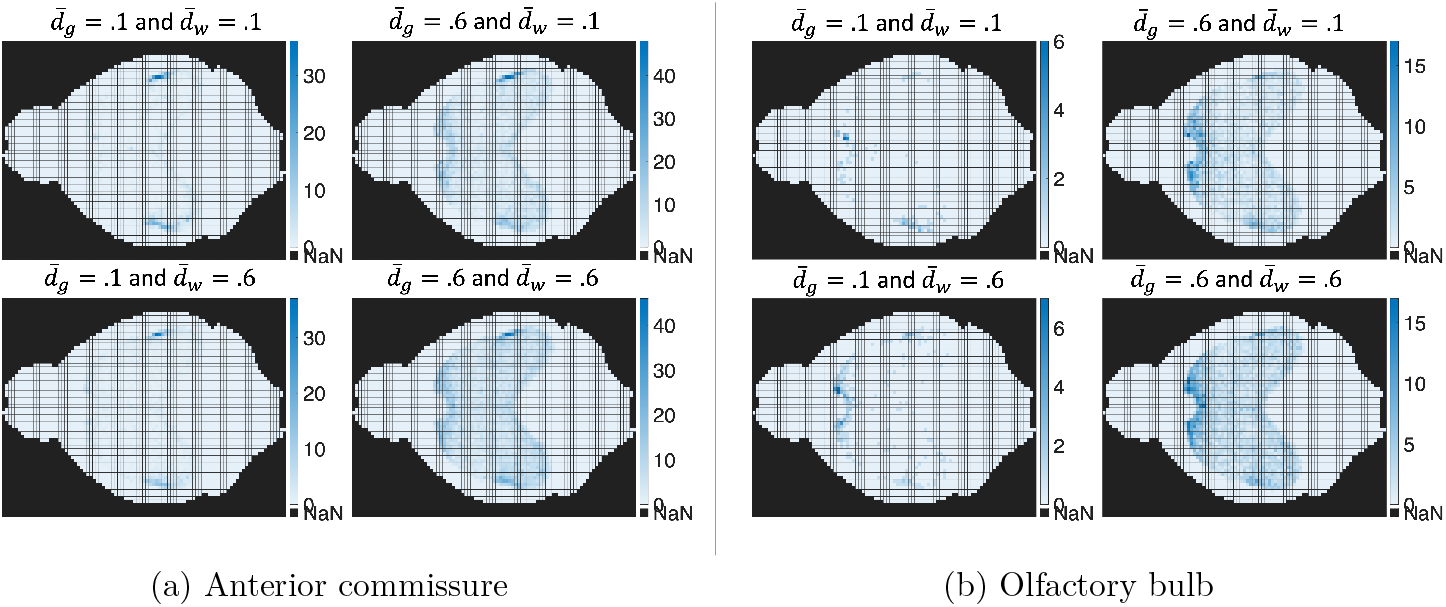
Heat maps showing NSC injection locations for cells reaching the target regions, (a) anterior commissure and (b) olfactory bulb. Quantities show the maximum count for a vertical slice in the *X/Y* region.

In Figure 7(a), we observed that the injection locations that most consistently led to the anterior commissure occur along the lateral edges of the corpus callosum. When the migration step sizes are small, 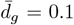 and 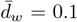, this is one of the only locations that lead to the anterior commissure. However, as we increase both directed and random migration step sizes, more locations in the brain show up including all along the edges of the corpus callosum and in particular the front of the corpus callosum. In general, with larger step sizes more cells end up reaching the anterior commissure partly due to random motion. However, we note that persistent region across all parameter combinations is the lateral edges. For cells that reached the olfactory bulb shown in Figure 7(b), a similar behavior is observed, but the persistent region in this case occurs in the front part of the corpus callosum.

To identify frequent pathways to the two target regions, we perform path clustering on the trajectories that made it to the anterior commissure and the olfactory bulb. This is done using *k*-medoids, where the distance between two trajectories is calculated as follows: first, at every 100th time step of the trajectory, the coordinates of the cell are binned by its position in the GQI image. Then a unique list of bins is created for each trajectory. Finally, the distance between two trajectories is calculated depending on the number of bins they have in common. In this way, if two trajectories share the exact same list of unique bins, the distance is zero.

In Figure 8, we plot the center of the *k*-medoids clustering with *k* = 7. Here, we note some key findings. Most of the representative trajectories (medoids) to the anterior commissure start on the lateral sides of the corpus callosum and travel down to the temporal limb of the anterior commissure. In fact, 5 of the 7 trajectories shown in Figure 8(a) follow the white matter fibers in this region. In addition, we see two other paths, one which passes from the corpus callosum to the fornix, and finally down to the anterior commissure (red), and one which passes from the front of the corpus callosum down to the temporal limbs of the anterior commissure (green). In contrast, for trajectories that lead to the olfactory bulb, shown in Figure 8(b), we see that cells either pass through the anterior commissure through the temporal and olfactory limb, before migrating to the olfactory bulb (green and purple), or they follow a path down from the front of the corpus callosum toward the olfactory limb of the anterior comissure (yellow and orange). Lastly, in red, we see that the final medoid represents randomized motion through grey matter which ends in the olfactory bulb. We note that in the experiments described in section 3, the NSCs are injected near the right side of the corpus callosum. Among the combined simulated paths leading to the anterior commissure and the olfactory bulb obtained, the only injection site at the front of the corpus callosum is the one that follows a trajectory through the olfactory limb before migrating toward these regions.

**Figure 8.**
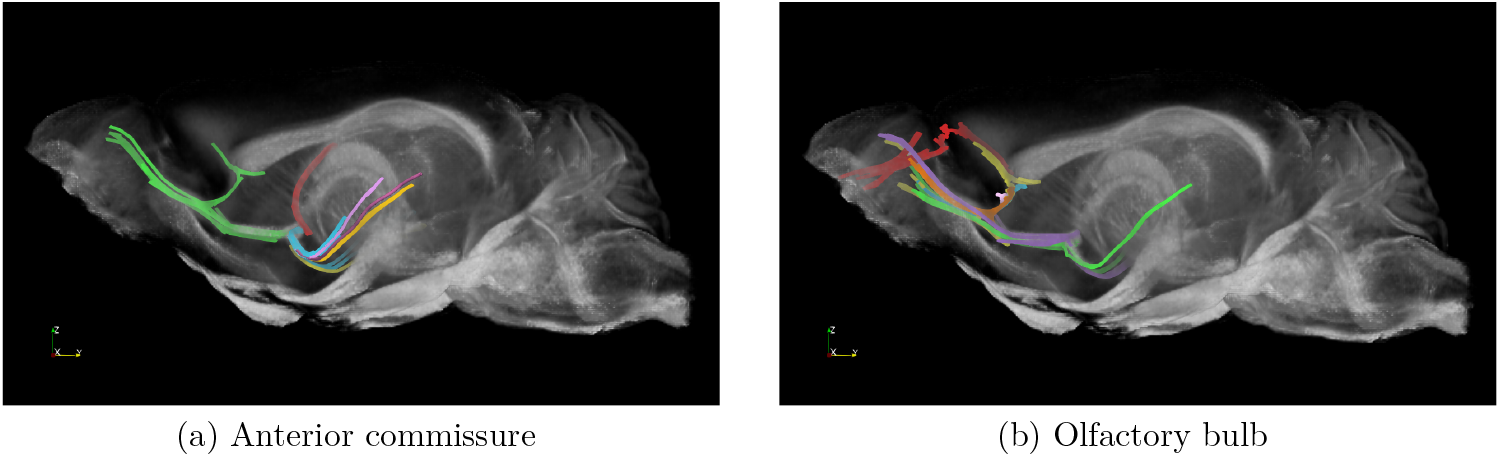
Predicted migration pathways of NSC trajectories reaching the target regions, (a) anterior commissure and (b) olfactory bulb. Collections of paths are identified with k-medoids clustering with *k* = 7 and shown in different colors.

### 4.4 Significance of Injection Location in Target Arrival Rates and Model Calibration

Using the estimated experimental injection location and corresponding experimental data, we calibrated the model parameters, 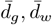, *K* based on the results of the sensitivity analysis (Figure 5). We estimate the parameter set that minimizes the sum of squared errors of migration distance and percentage in white matter at all time points. The set of optimal parameters are 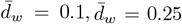, *K* = 0.18. See Figure S10 for the heat maps of the errors, which was narrowed down to 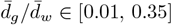 and *K* ∈ [0.22, 0.27] based on the error terms.

We then took the calibrated parameter set, 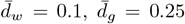, *K* = 0.18, and looked at the percentage of cells that arrive at the anterior commissure and olfactory bulb. Here we notice that the overall distribution of NSCs in the olfactory bulb is relatively small, with around 3.8% of the cells in the anterior commissure at 6 months and 7.7% of the cells being located in the olfactory bulb at 9 months. Since the experimental data is 7.2% and 19.9% respectively, we observed that the optimal parameters still did not capture the experimental results. This is because the calibration of 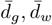, and *K* is effected by the injection location. To illustrate this, we took the optimal parameter set found above and run simulations in six alternate injection locations (2–6) which were around the most viable locations found in section 4.3. Note that cells are injected into a spherical region with radius 0.135 mm. Figure 9 shows where these regions are located, while Table 2 shows the results of each simulation. We see that the original location (1) and adjacent locations (2, 6) have the highest percentage of cells in the anterior commissure and olfactory bulb, while all the other locations show significantly lower percentages here. Moreover, the error from the median distance and percentage in white matter also varied significantly.

**Table 1:** Model parameters and their descriptions. *Parameter values are obtained directly from GQI reconstruction. Note our imaging data is stored on a 3D-matrix with Δ*x* = Δ*y* = Δ*z* = 45*µm* and 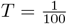 day.

| Parameter | Biological Meaning | Value |
| --- | --- | --- |
| $QA_i$ | Level of diffusion along the $i$th fiber resolved from GQI | * |
| $dir_i$ | The $i$th fiber direction resolved from GQI | * |
| MD | Mean diffusivity | * |
| $n$ | Hill coefficient | 7 |
| $K$ | Level of anisotropy in which motion is half directed and half random | .18 |
| $d_g$ | Maximum step size of NSC migration on grey matter | .25 |
| $d_w$ | Maximum step size of NSC migration in white matter | .1 |
| $\alpha_w$ | Stochasticity in migration speed parameter 1 | 1 |
| $\beta_w$ | Stochasticity in migration speed parameter 2 | 1 |

**Table 2:** Calibration error and percentages of NSCs arrived in anterior commissure (AC) and olfactory bulb (Olfac) at 6 and 9 months for simulations run at different injection locations (1–7) shown in Figure 9.

| Location | Exp. Error | AC 6 Month% | AC 9 Month% | Olfac 6 Month% | Olfac 9 Month% |
| --- | --- | --- | --- | --- | --- |
| 1 | 0.142 | 3.6% | 6.7% | 3.8% | 7.8% |
| 2 | 0.308 | 3.5% | 7.5% | 3.4% | 8.0% |
| 3 | 1.050 | 0.7% | 1.0% | 1.0% | 1.5% |
| 4 | 6.774 | 0.4% | 0.2% | 0.7% | 0.3% |
| 5 | 1.023 | 0.3% | 0.1% | 0.5% | 0.1% |
| 6 | 5.401 | 1.6% | 3.4% | 1.7% | 3.9% |
| 7 | 0.632 | 0.2% | 0.1% | 0.3% | 0.2% |

**Figure 9.**
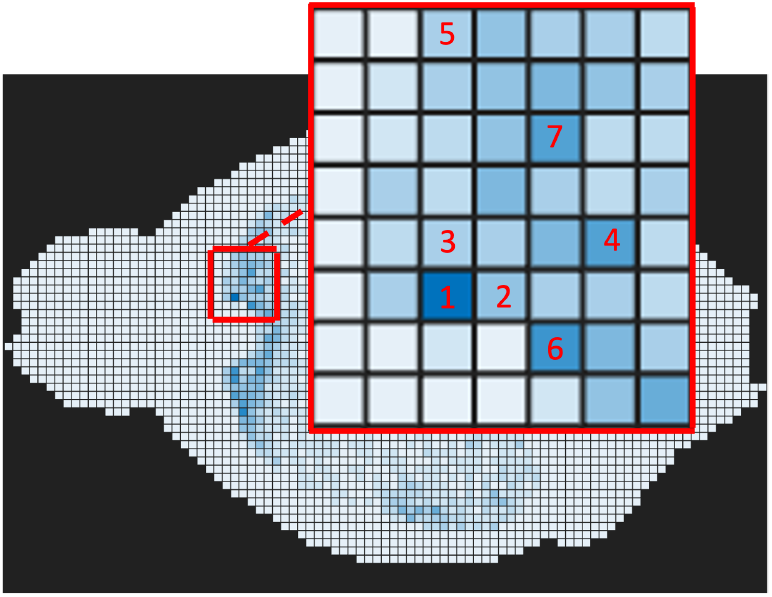
Heat map showing the injection sites for NSCs reaching the olfactory bulb at any point during the simulated 9 months and the selected seven injection locations (1–7) that were tested. The heat map shown is for simulations with parameters 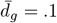 and 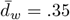

Since small changes to the injection locations can result in large changes on the measured arrival percentages, we hypothesized that differences in the injection location could explain some of the variation we see between mice in the experimental data. For example, the estimates we found for the number of cells that made it to the olfactory bulb for each individual mouse were 33.3%, 20.3%, 4.2%. This illustrates differences observed between the mice. Our model similarly was able to demonstrate large differences in the percentage of cells that are able to make it to given regions; however, less percentage of cells arrive at olfactory bulb in our model simulation compared to the experimental data. Hence we conclude that this behavior is either a product of injecting into a very specific location, or that tissue anisotropy alone is not adequate enough to explain the experimental results.

## 5 Discussion

In this paper we present a probabilistic, agent-based model of neural stem cell migration in naïve mouse brain that accounts for multiple crossing white matter fibers. Using high-resolution GQI reconstruction, we are able to more accurately simulate the migration of NSCs along secondary and tertiary fibers within the brain. We demonstrate this ability by simulating the migration of 100 NSCs in a region of dense fiber crossings and find that the cells are better able to traverse the resolved fiber directions when compared to a model which only uses DTI data.

We conclude that both the migration distance and the percentage of NSCs in white matter is an important quantity for calibrating the model parameters. From the sensitivity analysis, we find that the most influential parameters on the median distance and percent in white matter were 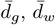, and *K*, while *α, β*, and *n* had minimal effect. The step size parameters 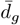 and 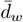 are positively correlated with the migration distance, however, very low 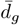 and 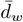 values yield an opposite correlation for 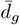. From this we hypothesize that for low step sizes the decrease in the median distance due to increasing 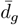 is likely due to an increased accumulation of cells on the local white matter tracts which traps cells and decreases their ability to travel to other regions of the brain. We show that the ratio 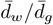 is inversely related to the change in percentage in white matter demonstrating the usefulness of the temporal changes in white matter percentages for model calibration. However, we note that our work utilizes experimental data collected at three time points - 3, 6, and 9 months - and suggest that additional temporal data not only would enhance the parameter calibration of this model, but might also uncover additional mechanisms underlying NSC migration in the brain.

We also performed path clustering on NSCs injected uniformly throughout the corpus callosum and found that there was only one main white matter path that led to the anterior commissure and the olfactory bulb in the right anterior region of the corpus callosum. It is the one that follows a trajectory through the olfactory limb before migrating toward these regions. Therefore, we predict that the NSC injection site in the experiments would have overlapped with this identified location. We further investigated the injection location and found that this had a significant effect on the migration path and overall distribution of the cells. For instance, the percentage of NSCs that arrived at the anterior commissure was significantly different by an order of magnitude with only a couple of hundred *µm* difference in the injection location. Finally, we emphasize that our model can identify potential injection sites and likely migration paths to any target location, offering a promising approach to improve the efficacy of NSC treatments.

Our model is calibrated to the experimental data for the migration of NSCs over 9 months. We found that the model is able to fit the quantitative data for the experimental median distance from injection site and percentage in white matter; however, it struggles to reproduce the distribution of NSCs distribution in the olfactory bulb that was observed experimentally in some mice at 9 months. We emphasize that the injection location of NSCs has a significant effect on the migration path and overall distribution of the cells, suggesting that either this qualitative behavior is a product of injections being done in a specific location or other forces not included in the model. For example, blood vessels have been shown to play a role in neuronal migration both in neonatal and adult rodents. Notably, some studies have shown that neuroblasts derived in the sub-ventricular zone of neonatal mice are able to travel to the cortex through the corpus callosum by use of the vasculature network [25]. In addition, VEGF helps to promote the growth of blood vessels that run parallel to the rostral migratory stream [9]. These vessels in adult mice, may provide neuroblasts with a scaffold as well as excrete various chemical signals that promote migration [41, 47]. Due to findings like these, recent studies have begun to model the migration of neuroblasts from the subventricular zone to the olfactory bulb as a chemotactic process [2]. For a more in depth review of these topics see the following review [16]. Because of findings like this, chemotaxis as well as cell-cell adhesion may not only be an important driving force in NSC migration towards injury, but may also play a key role in migration in naïve brain as well.

Our work emphasizes the importance of advanced imaging techniques, such as GQI, that can characterize individual brain tissue structures. This can be followed by a personalized injection location precisely determined to improve the efficacy of NSC treatments for specific target locations.

## Declarations

### Data/code availability

Raw data of the mouse brain DTI is supplied by Duke Center for In Vivo Microscopy (CIVM) [45] and reconstruction was done in DSI studio [1]. The NSC experimental data is supplied by Dr. Margarita Gutova’s group available in [38]. The code will be available in https://github.com/AustinHansen82 upon publication.

## Acknowledgment

Thanks to James J. Cook from Duke CIVM for technical assistance.

## Supplementary Information

### S1 Parameter Exploration

In this section we explore the relationship between the model parameters 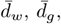, *K, n, α*, and *β*. We expand on the observations concluded from the Partial Rank Corelation Coefficients (PRCC) shown in section 4.2, by running three additional sets of 100 simulations each. In each set, 2 parameters are varied over a stated range, and all other parameters remain constant. We then plot the effect on the median distance from injection site, as well as the percentage of cells on white matter over the course of the simulated 9 months. For all simulations, 10,000 cells were injected into a spherical region of radius .135*mm* in the right anterior of the corpus callosum, so as to approximate to the experimental injection site in [38].

#### S1.1 Step Size Analysis

**Figure S1:**
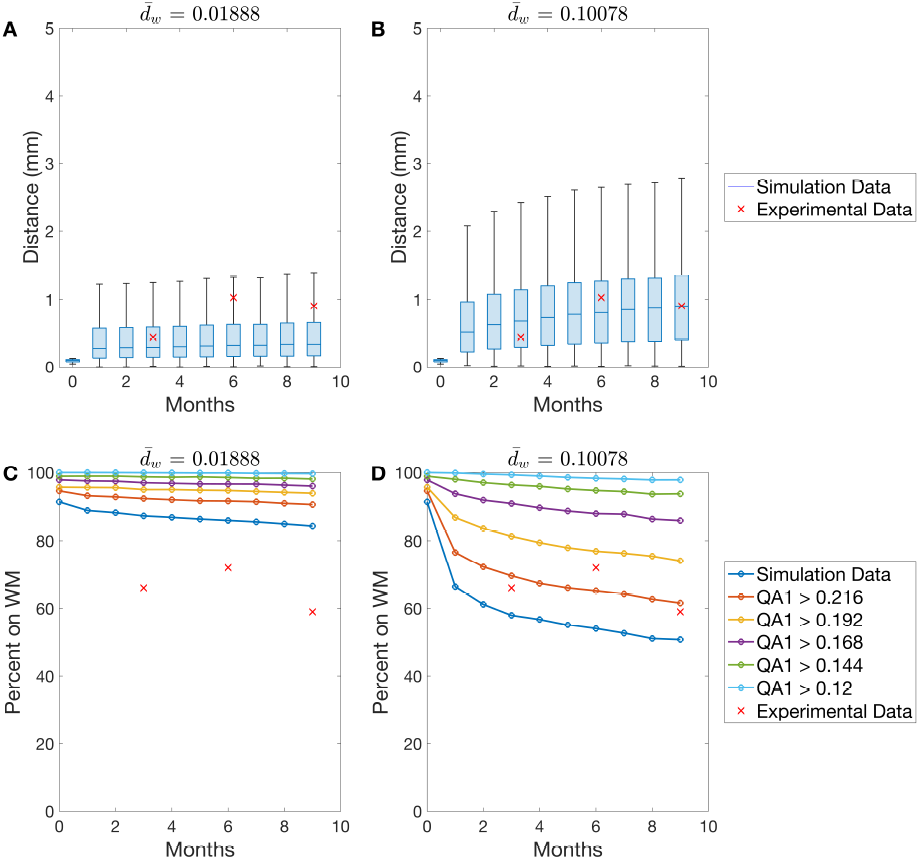
Box plots of the distances from injection site as well as the percent in white matter across the 9 months of simulated NSC migration. Here 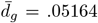 is fixed. A and C show 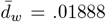. While B and D show 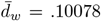. We see as 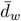 increases the median distance increases while the percent in white matter decreases.

We first turn our attention to the effects of 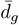 and 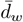. As shown through the PRCCs, 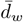 is positively correlated to the median distance and is inversely related to the percent in white matter. This is further illustrated in Figure S1. Here we compare 2 simulations where 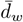 is varied between .01888 and .10078, while *n* = 7, *K* = .24, *α* = *β* = 1, and 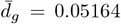. Figures S1A and S1B show the box plots for the start of each simulated month, as well as the experimental distances at months 3, 6, and 9 shown in red. We clearly see an increase in the median distance from the injection site as we increase 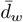. In order to quantify the percentage of NSCs on white matter, we consider a NSC at location **x** to be on white matter if *QA*_1_(**X**) > *K* and grey matter otherwise. Here **X** is the nearest lattice point to the point **x** which may be off lattice. This designates every point in the brain with a primary QA value greater than *K* as white matter. Since QA is relatively smooth we also plot the interface around the white matter tracts by considering **x** to be on white matter when 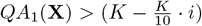 for *i* ∈ [0, 1, 2, 3, 4, 5]. Looking at Figures S1C and S1D, we can again clearly see that increasing 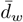 leads to a decrease in the percentage on white matter.

**Figure S2:**
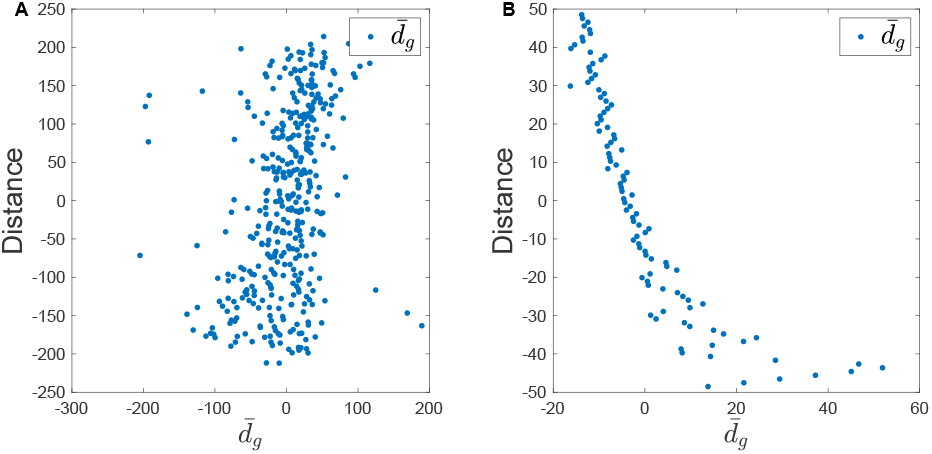
Plot showing the residuals 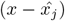 and (*y*_*j*_ −*ŷ*) where *x* and *y* are the rank transformed 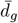 values and 9 month median distance respectively and 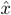, *ŷ* are from the linear regression models. A shows the residuals for sensitivity analysis run across the entire relevant parameter space while B shows the residuals for simulations with 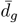 ∈ [.0025, .15].

**Figure S3:**
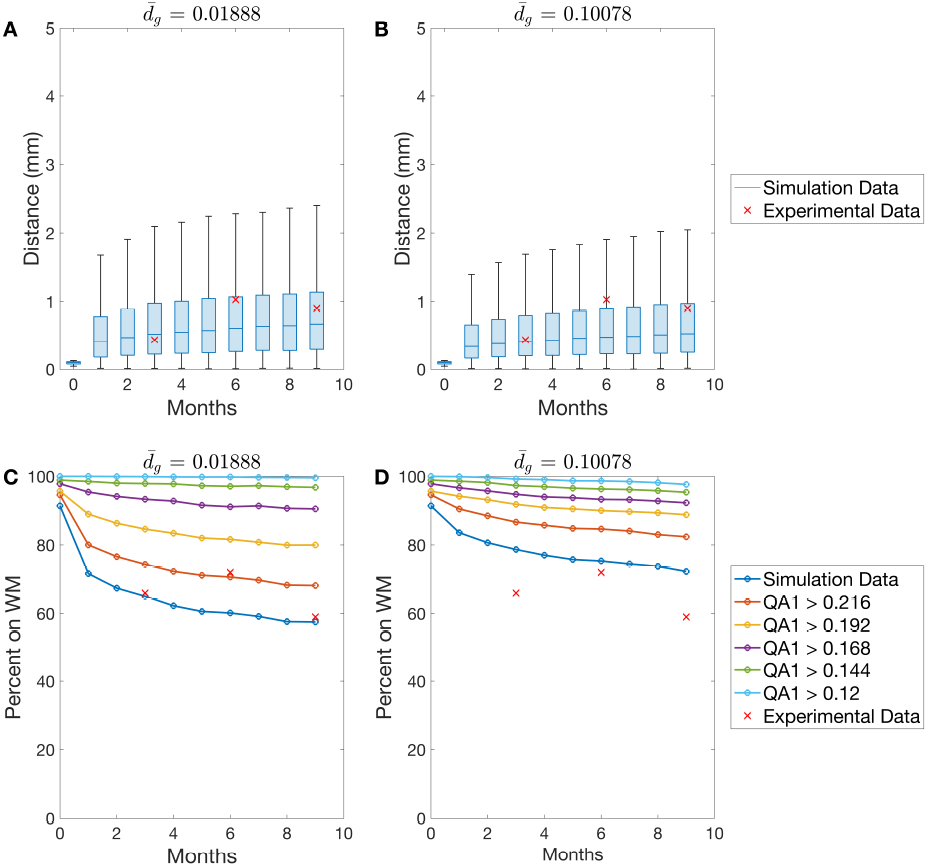
Box plots for the distances from injection site as well as the percent in white matter across the 9 months of simulated NSC migration. Here 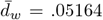 is fixed. A and C show 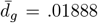. While B and D show 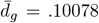. We see as 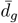 increases the median distance decreases while the percent in white matter increases.

In contrast, 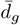 is positively related to both the median distance and the percent in white matter. The positive correlation between both step sizes and the median distance from the injection site is intuitive, as the larger the step sizes, the faster the cells travel in the brain. However, as mentioned in the main text, the effect of 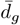 on the median distance is partially dependent on the range of step sizes considered when calculating the PRCC. In Figure S2 we plot the residuals 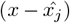 and (*y*_*j*_ − *ŷ*) where *x* and *y* are the rank transformed 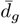 values and 9 month median distances respectively, and 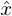, *ŷ* are from the linear regression models. In Figure S2A we depict the residuals from the full Latin hyper-cube sampling of 400 simulations with parameter values sampled from the following ranges: 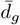 and 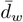, *K* ∈ [0, .5], *n* ∈ [1, 30] and *α, β* ∈ [1, 100]. In Figure S2B, the depicted residuals come from a set of 100 simulations where 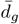 and 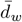 are selected from the range [.0025 : .01638 : .14992] while *n* = 7, *K* = .24, *α* = *β* = 1 are held constant.

In comparing the residuals we notice that for smaller 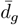 and 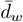 values, the correlation is actually negative. We illustrate this in Figure S3. Here 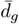 is varied between .01888 and .10078 while *n* = 7, *K* = .24, *α* = *β* = 1, and 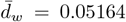. Figures S3A and S3B show that as 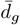 increases, the median distance decreases. This is true for all 9 simulated months. Furthermore, in Figures S3C and S3D, we can see the inverse relationship between the percent in white matter and 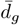. As discussed in section 4.2, the accumulation of the NSC’s in white or grey matter is governed by the ratio 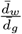. Thus, an increase in 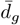 actually increases the tendency of the cells to accumulate on white matter. We hypothesize that when the step sizes are small, the increased accumulation on white matter may lead to a disproportionate decrease in the ability of cells to leave their local white matter tracts. This could lead the NSCs to become trapped in one area of the brain, ultimately lowering the median distance from the injection site. However, when the step sizes are larger, this increased accumulation may not be as impactful, as the cells will move fast enough to traverse the grey matter areas of the brain before finding other white matter fibers to follow.

#### S1.2 Hill Parameters Analysis

In this section we illustrate the effects of the parameters *K* and *n* on the median distances from the injection site and percent in white matter. The simulations shown in Figure S4 are taken from a set of 100 simulations with *K* ∈ [.1 : .04 : .46] and *n* ∈ [3 : 2 : 21]. The rest of the parameters are fixed with 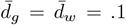 and *α* = *β* = 1. The first observation we make is that as we increase *K*, the median distance from the injection site decreases. Since *K* is the point where half of the cell’s velocity comes from directed motion and half is random, this implies that as cells move with more randomized motion, the lower the median distance from injection site will be. Since *K* also directly controls the QA values that are considered white matter, we see that increasing *K* decreases the percent in white matter as expected. This happens because less regions in the brain are considered as white matter with increased *K*.

**Figure S4:**
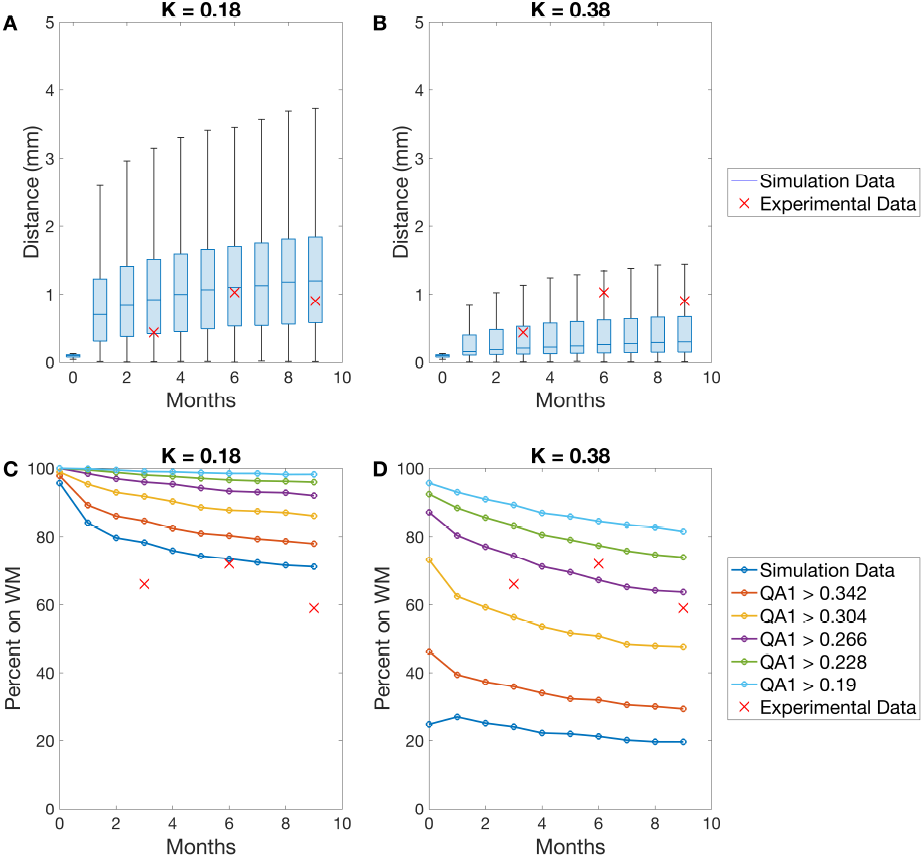
Box plots for the distances from injection site as well as the percent in white matter across the 9 months of simulated NSC migration. Here *n* = 7 is fixed. A and C show *K* = .18. While B and D show *K* = .38. In We see as *K* increases the median distance as well as percent in white matter decreases.

**Figure S5:**
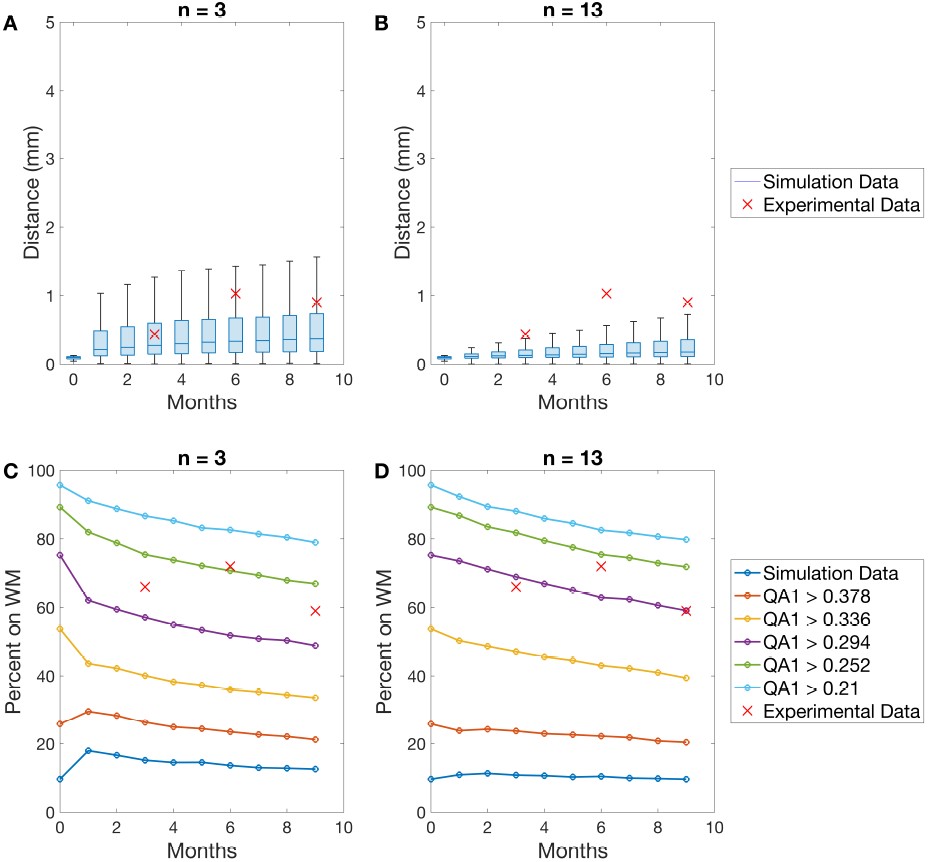
Box plots for the distances from injection site as well as the percent in white matter across the 9 months of simulated NSC migration. Here *K* = .42 is fixed. A and C show *n* = 3. While B and D show *n* = 13. We see as *n* increases the median distance as well as the variance also decreases. The percent in white matter is relatively unaffected.

**Figure S6:**
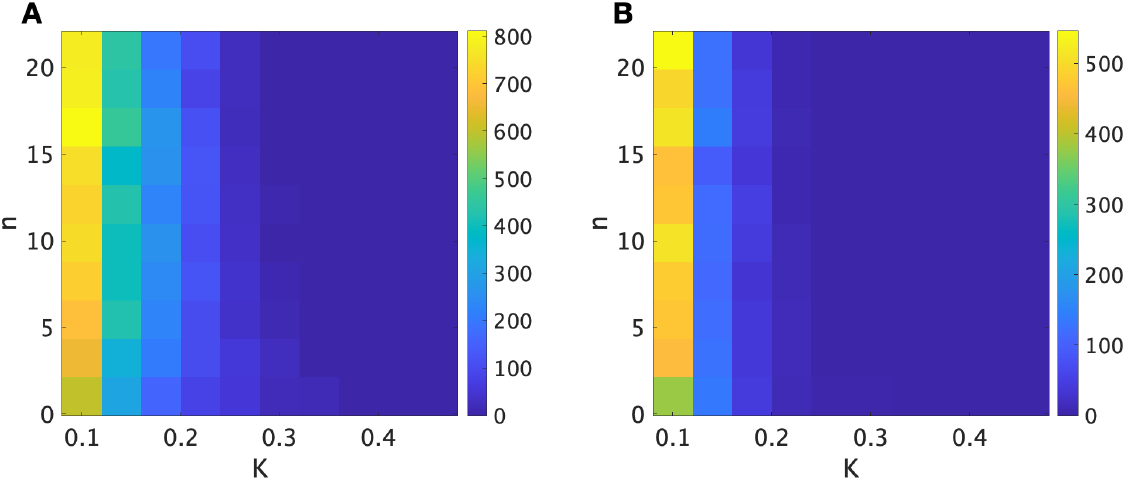
Heat map showing the number of cells that are within the anterior commissure (A) and the olfactory bulb (B) at any point in the simulation. We see that for lower *K* values the great amount of cells make it to the anterior commissure and corpus callosum respectively.

Now, considering a fixed *K* and a variable *n*, we find that the parameter *n* does not make much difference. The only notable trend shows up when we have a large fixed value of *K* > .42, which is depicted in Figure S5. Here we see that when *n* increases, the median distance decreases. This is because cells are unable to leave the local white matter tract. Since *K* is large, most parts of the brain are considered grey matter and thus an increase in *n* causes this grey matter motion to be completely random. This causes it to become increasingly less likely that cells are able to travel across large potions of grey matter and find other white matter tracts to explore.

Finally, if we consider how many cells make it to the anterior commissure and olfactory bulb at any point in the simulation, we note one more interesting trend. In Figure S6 we plot heat maps showing the number of cells that made it to the anterior commissure and the olfactory bulb at any point in the simulation. Again, *n* does not play a big role in either of these behaviors; however, we see that as *K* decreases the amount of cells that make it to these structures increases. This suggests that as more motion comes from the tissue structure, the more migration we get to the anterior commissure and the olfactory bulb.

#### S1.3 Stochasticity Analysis

Finally, we illustrate the effects of varying the stochasticity parameters *α* and *β*. Here we fix *K* = .24, *n* = 7, 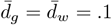. We point out that for a given beta distribution, *X* ∼ Beta(*α, β*), as *β* increases, the distribution skews towards the left, and thus more cells have smaller step sizes. Inversely as *α* increases, the distribution skews toward the right, and more cells have a larger step size. In order to isolate only the effects of stochasticity we compare the results for the extreme case when *X* is uniform with *α* = *β* = 1 and when *X*(*x*) = .5 for all *x*. Note that both have the same expected value of .5. Furthermore, in order to eliminate the noise from starting locations, we initialize the NSC’s at a single location within the brain. In Figure S7 we see that there is little effect on the median distance and percent in white matter when the expected value is fixed. However, this stochasticity parameter does effect the number of outliers. We see that in A there are considerably more outliers for the uniform distribution in comparison to the fixed value in B. This suggests that the expected value is the biggest contributor to the median distance and percent in white matter.

**Figure S7:**
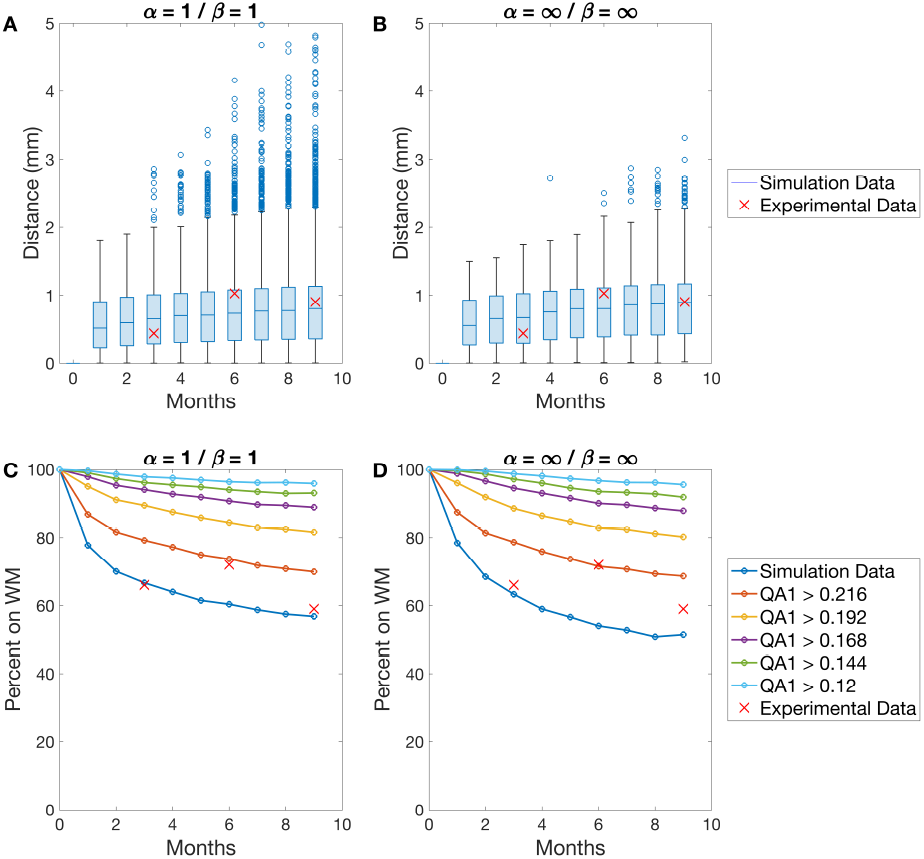
Box plots for the distances from injection site as well as the percent in white matter across the 9 months of simulated NSC migration. A and C show a uniform distribution with *α* = *β* = 1. B and D show the case when *X*(*x*) = .5 for all *x*. We see that with the stochasticity parameters only effects the outliers and not the median distances.

We can again look at the heat map for the number of cells that made it to the anterior commissure and olfactory bulb coming from 100 simulation with *α, β* ∈ [1, 10]. We see in Figure S8 that the number of cells that reach the anterior commissure and olfactory bulb is highest when *β* is low and *α* is large. Since varying *α, β* changes the expected value of the distributions, the average velocities of the NSC will have the highest velocity in this region. Therefore, this result can be interpreted as just showing that increased cell velocity increases the probability that the NSCs arrive at the anterior commissure and olfactory bulb.

**Figure S8:**
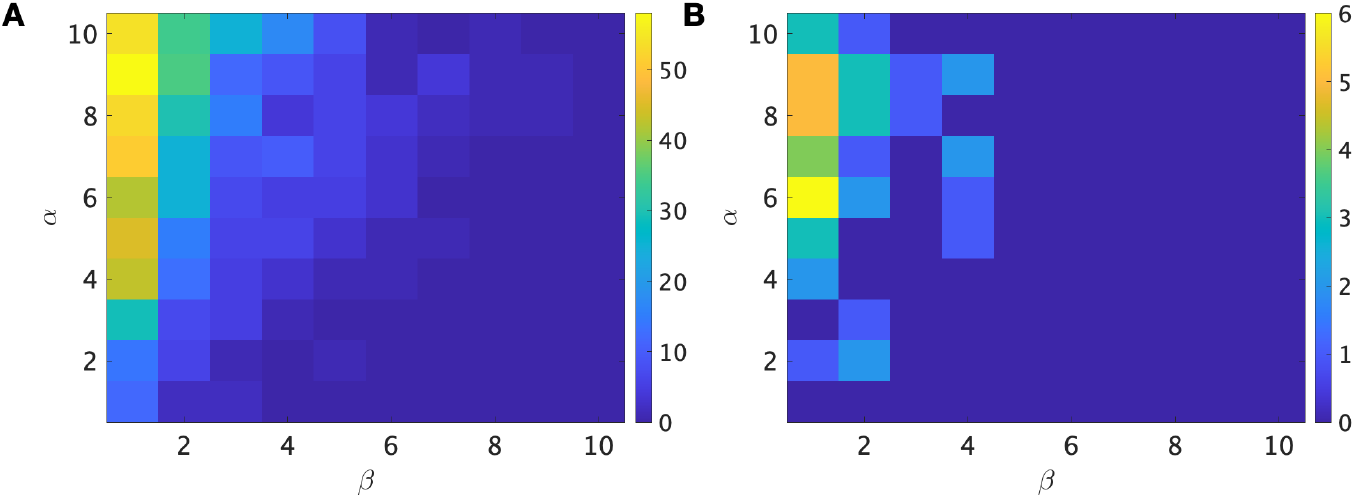
Heat map showing the number of cells that are within the anterior commissure (A) and the olfactory bulb (B) at any point in the simulation. We see that the lower *β* and higher *α* values the great amount of cells make it to the anterior commissure and corpus callosum respectively.

### S2 Experimental Calibration

In this section we explain the details of the model calibration. From the above parameter analysis we can conclude that the parameters 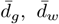, and *K* have the largest influence on the simulated trajectories. Since *α* and *β* only effect the amount of outliers, we fix *α* = *β* = 1. Furthermore, we set *n* = 7. We make this selection since for *K* ≈ .25, a value of *n* = 7 yields *w*_*QA*_ ∈ [.2, .8] for around 20% of the primary QA values. This means 20% of locations in the brain will lead to motion which is a combination of directed and random, such that each is weighted between 20% − 80%, while the rest of the QA values will lead to motion which is either mostly random, or directed along the white matter tracts. To depict the relationship between *K, n*, and the values *QA* in the brain, we plot the hill function for *n* = 7 and *K* ∈ [.1, .46]. In addition, for each function we show the percentage of cells within the brain that have a primary *QA* such that *w*_*QA*_ ∈ [.2, .8]. Here we can see that given *n* = 7, the maximum number of cells in this range occurs for *K* = .14 and is a unimodal distribution around this maximum.

**Figure S9:**
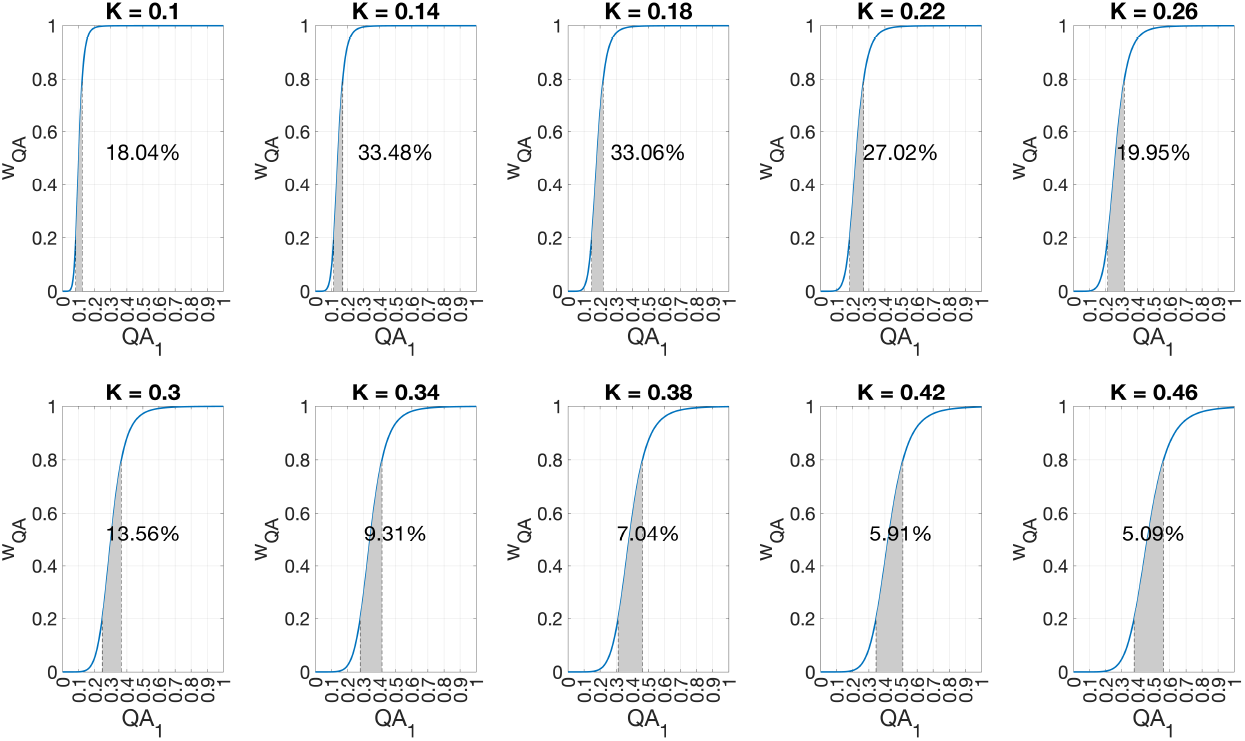
Plots showing the Hill function for *n* = 7 and *K* ∈ [.1, .46]. the shaded area represents the portion of the function in which *w*_*QA*_ ∈ [.2, .8]. The percentage stated shows the percentage of cells in the brain such that *QA*_1_ lies in this region

We then ran a host of simulations varying 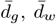, and *K* and calculated the sum of the squared error term between the simulated and experimental values at 3,6,9 months. The following set of heat maps shows a narrowed-down range for this analysis. The simulation outlined in green represents the simulation with the smallest error term showing a value of .132.

**Figure S10:**
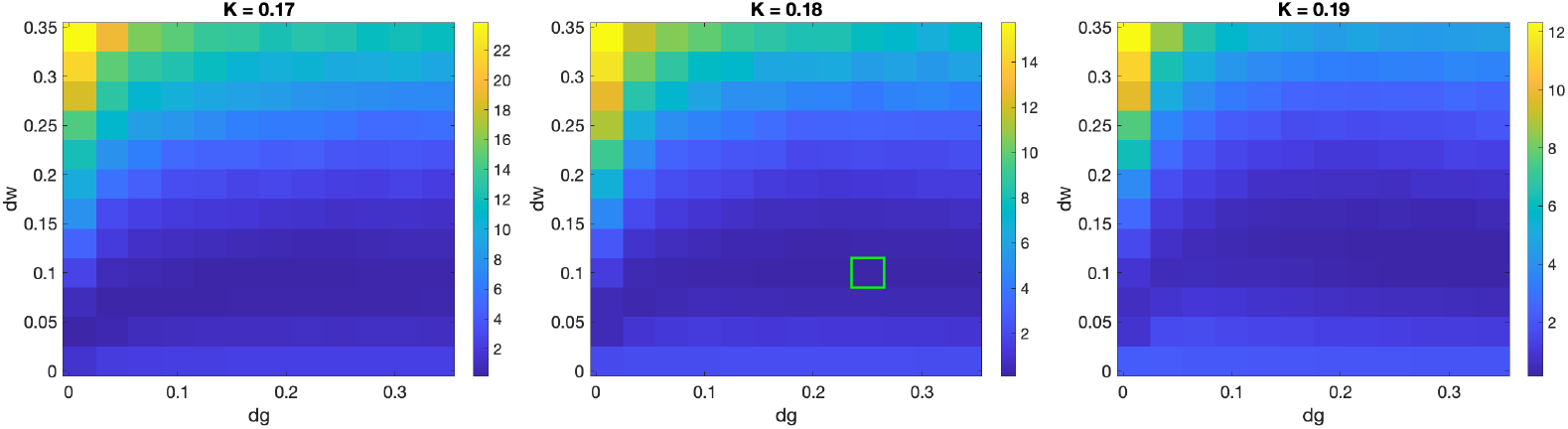
Heat showing the sum of squared error between the simulated and experimental median distances from injection and percentage on white matter at 3,6,9 months. The green highlight show the simulations with the best error which occurs with parameter values 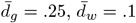, and *K* = .18

